# The multiple lncRNAs encoding *hsr*ω gene is essential for oogenesis in *Drosophila*

**DOI:** 10.1101/2022.12.24.521879

**Authors:** Rima Saha, Subhash C. Lakhotia

**Author notes:** E-mail : Rima Saha, S. C. Lakhotia.

## Abstract

In the background of limited studies on noncoding RNAs in *Drosophila* oogenesis, we show developmentally active *hsr*ω lncRNA gene to be essential in oogenesis and ovulation. The near-null *hsr*ω*^66^* females, and ovaries with down- or up-regulated *hsr*ω display varyingly perturbed oogenesis including fewer ovarioles, high apoptosis, poor actin nuclear-cage (stage 10), low Cut levels in late chambers and, finally ovulation block. Restoration of normal oogenesis following targeted expression of *hsr*ω-RH transcript in *hsr*ω*^66^* confirmed *hsr*ω mis-function to underlie these defects. Genetic interaction studies showed varying modulation of ovarian defects following mis-expression of Cut, and TBPH/TDP-43 or Caz/dFUS hnRNPs by altered *hsr*ω transcript levels. Dietary supplement of ecdysone to *hsr*ω*^66^* females, which have reduced ecdysone titer, substantially restored normal oogenesis. Our results show for the first time that the multiple lncRNA producing *hsr*ω gene, which interacts with diverse hnRNPs and other regulatory molecules, As expected of a gene with key roles in dynamics of various hnRNPs, interactions between down or upregulated *hsr*ω transcripts and various oogenesis regulators are not linear.

**Summary Statement:** The multiple lncRNA producing *hsr*ω gene critically impacts *Drosophila* oogenesis at multiple steps through intra- and inter-organ signaling.

## Introduction

Oogenesis is a complex process involving multiple regulatory pathways working through intra- and inter-organ signaling (Doherty et al., 2022; Merkle et al., 2020; Spradling et al., 2022). Despite the increasing attention received by diverse long noncoding RNAs (lncRNAs), relatively fewer studies have examined their roles in oogenesis. While several lncRNAs are shown to have roles in mammalian oogenesis (Tu et al., 2020), only the *oskar* cncRNA (Sampath and Ephrussi, 2016) is known to be important in *Drosophila* oogenesis (Song and Zhou, 2020). Earlier studies in our lab showed high expression of developmentally active and cell stress inducible *hsr*ω lncRNA gene (Lakhotia, 2011) in ovarian nurse and follicle cells in *Drosophila melanogaster* (Mutsuddi and Lakhotia, 1995). Female fecundity was low following targeted reduction of *hsr*ω transcripts (Mallik and Lakhotia, 2011). The *hsr*ω lncRNA gene produces 7 transcripts using two transcription start and four termination sites and differential splicing of its single intron (Sahu et al., 2020).

We now examined oogenesis in *hsr*ω*^66^*, a near-null allele carrying a 1.6 kb deletion spanning the proximal part of promoter and first 9bp after the second (major) transcription start site (Johnson et al., 2011) and which produces only traces of different *hsr*ω transcripts (Ray et al., 2019a). The *hsr*ω*^66^* homozygous individuals are unhealthy (Johnson et al., 2011) with reduced fecundity (Mallik and Lakhotia, 2011). In present study, the *hsr*ω*^66^* ovaries, and those with targeted down- or up-regulation of *hsr*ω transcripts were found to be poorly developed with defective oogenesis and ovulation block. Targeted over-expression of *hsr*ω *RH*, one of the transcripts of this gene, in ovarian follicle cells significantly reversed the oogenesis defects associated with *hsr*ω*^66^* homozygosity. Genetic interaction studies involving targeted co-down- or up-regulation of *hsr*ω transcripts and Notch, Cut, Caz/dFus or TBPH/TDP-43 during oogenesis revealed non-linear consequences of altered levels of one or more of these interactors, as expected of a gene producing multiple lncRNAs that affect a variety of regulatory events in cells (Lakhotia, 2011; Ray et al., 2019a; Ray et al., 2019b; Singh and Lakhotia, 2015). Remarkably, providing dietary ecdysone to *hsr*ω*^66^* flies also significantly improved oogenesis. The present results thus show that the *hsr*ω lncRNAs modulate multiple intra- and inter-organ regulatory events to ensure successful completion of the complex process of oogenesis.

## Results

### The *hsr*ω*^66^* females show reduced life span, poor fecundity, and ovulation block

The *hsr*ω*^66^* homozygotes are near-null for all the *hsr*ω transcripts (Johnson et al., 2011; Ray et al., 2019a). The *w^1118^* flies, which show normal lifespan and fecundity comparable to the wild type flies, were used as control in all cases since all the examined genotypes carried *w^1118^* or other *white eye* mutant allele.

None of the *hsr*ω*^66^* homozygous females, when kept with *hsr*ω*^66^* homozygous males, survived beyond 33 days after emergence, although nearly 90% the *w^1118^* females, maintained with *w^1118^* males or with *hsr*ω*^66^* homozygous males, survived (Fig. 1A). Interestingly, keeping *hsr*ω*^66^* homozygous virgins with *w^1118^* males further reduced their viability these females did not survive beyond 18 days (Fig. 1A).

Fecundity assay of *hsr*ω*^66^* homozygous females, measured as mean numbers of eggs laid by one female per day till 14 days after emergence, revealed that compared to *w^1118^* females, *hsr*ω*^66^* females laid fewer eggs (Fig. 1B). Interestingly, keeping *hsr*ω*^66^* virgin females with *w^1118^* males further reduced the mean numbers of eggs laid/day but keeping *hsr*ω*^66^* males with *w^1118^* virgins did not affect their egg-laying capacity (Fig. 1B). Oviposition by *hsr*ω*^66^* females was also delayed as they laid eggs only after 2 days post-emergence.

We examined *w^1118^* (control) and *hsr*ω*^66^* ovaries on different days (20 ovaries on each day) since eclosion (day 0). The ovaries were significantly smaller on day 0 in *hsr*ω*^66^* homozygotes than in same age controls (Fig. 1C, D). Unlike the presence of stage 7-8 egg chambers in *w^1118^* ovaries on day 0 (Fig. 1C), egg chambers beyond the stage 5-6were not present in same age *hsr*ω*^66^* ovaries (Fig. 1D). Unlike the presence of mature follicles ready for ovulation in 1-day old *w^1118^* females (Fig. 1E), same age *hsr*ω*^66^* homozygous ovaries showed only a marginal increase in ovary size but without mature follicles (Fig. 1F). In agreement with the beginning of oviposition by *hsr*ω*^66^* females on day 3 (Fig. 1B), *hsr*ω*^66^* ovaries showed presence of some mature follicles on day 3 (Fig. 1H), although the overall ovary size remained smaller than corresponding *w^1118^* ovaries (Fig. 1G). Compared to 6 and 8 days old *w^1118^* ovaries (Fig. 1I, K, M) similar age *hsr*ω*^66^* females had fewer ovarioles (Fig. 1J, L, M) with rare mid- and late-stage egg chambers. Interestingly, post-6 days *hsr*ω*^66^* ovaries carried multiple unovulated mature follicles (Fig. 1N) with abnormal dorsal appendages (Fig. 1L), features characteristic of ovulation block (Deady et al., 2017).

As described in Supplementary Results and Fig. S2, *hsr*ω*^66^* ovarioles showed reduced Vasa staining and weak fusomes in germarium and early chambers.

Unless otherwise noted, ovaries from 6 days old females of different genotypes were used in all subsequent studies.

**Fig. 1.**
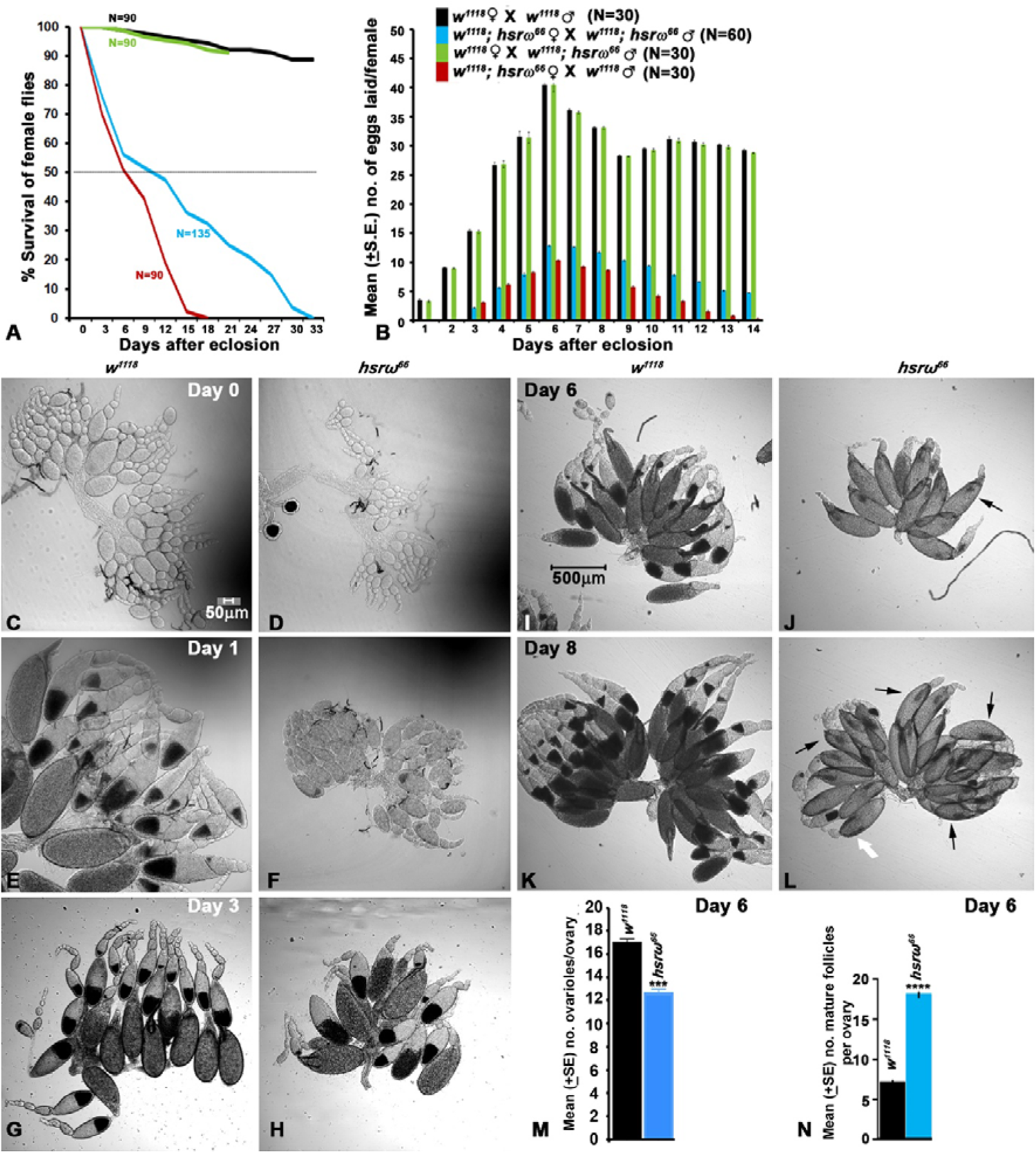
Near absence of *hsr*ω transcripts reduces lifespan and fecundity of *Drosophila* females, associated with poor ovarian growth, and ovulation block. **(A)** Lifespan (% survival (Y-axis) on different days (X-axis)) of *w^1118^* and *hsr*ω*^66^* homozygous females crossed with males of different genotypes (crosses noted on upper right corner in **B**). **(B)** Graphical presentation of the Mean (±S.E.) numbers of eggs laid (Y-axis) on different days after eclosion (X-axis) by *w^1118^* and *hsr*ω*^66^* homozygous females crossed with males of different genotypes (noted on upper right corner). N indicates the total numbers of flies examined (sum of 3 independent replicates) for each genotype. **(C-L)** Ovaries of *w^1118^* (**C, E, G, I,** and **K**), and *hsr*ω*^66^* (**D, F, H,** and **L**) females on day 0 (**C, D**), day 1 (**E, F**), day 3 (**G, H**), day 6 (**I, J**), and day 8 (**K, L**) after emergence; black arrows in **J** and **L** point to stacked mature follicles, while white arrow in **L** indicates abnormal dorsal appendages. (**M, N**) Mean numbers (± S.E.) of ovarioles per ovary (Y-axis, **M**) and mean numbers (± S.E.) of mature follicles per ovary (Y-axis, **N**) on day 6 in *w^1118^* and *hsr*ω*^66^* females, (N= 60 ovaries for each genotype). Scale bars in **C** (50 µm) and **I** (500µm) apply to **C-H** and **I-L,** respectively.

### Fasciclin III and F-actin reduced in *hsr*ω*^66^*

Distribution of Fasciclin III (Fas III), a follicle cell marker (Lee et al., 1997; Nystul and Spradling, 2010), in the germaria of *w^1118^* and *hsr*ω*^66^* (Fig. 2A-D) appeared significantly weaker in *hsr*ω*^66^* (Fig. 2I). Numbers and locations of different follicle cells in the few mid and late-stage follicles present in *hsr*ω*^66^* ovaries appeared similar to those in *w^1118^* ovaries (Fig. 2E, F). However, DAPI and Fas III staining revealed that in the mature *hsr*ω*^66^* egg chambers (stage 13 and 14), the body follicle cells showed disorganized linear columns (Fig. 2F, H) instead of the characteristic hexagonal pattern in *w^1118^* chambers (Fig. 2E, G). Further, the body follicle cell nuclei appeared larger in *hsr*ω*^66^* late chambers than in *w^1118^* chambers (Fig. 2G and H). The Fas III staining intensity in body follicle cells was also lower in *hsr*ω*^66^* chambers than in *w^1118^* (Fig. 2E-H and J).

Unlike the Fas III rich small corpus luteum at posterior end of mature follicles in *w^1118^* (Fig. 2K), the corpus lutea of multiple unovulated mature egg chambers in *hsr*ω*^66^* ovaries fused to make large Fas III positive bodies (Fig. 2L). The outer epithelial lining of proximal as well as distal parts of oviduct also showed reduced Fas III staining in *hsr*ω*^66^* (Fig. 2N, and P) than in *w^1118^* (Fig. 2M, and O).

Staining of *hsr*ω*^66^* and *w^1118^* ovaries with Phalloidin (Fig. 2Q-V) revealed similar F-actin distribution in mid-stage *w^1118^* (Fig. 2Q) and *hsr*ω*^66^* chambers (Fig. 2R) but unlike the robust F-actin nuclear cage in nurse cells of stage 10 *w^1118^* chambers (Fig. 2S), the *hsr*ω*^66^* chambers showed poorly formed nuclear cage (Fig. 2T). Mature *hsr*ω*^66^* chambers also displayed significantly reduced F-actin in the outer muscle layer (Fig. 2U, V).

**Fig. 2.**
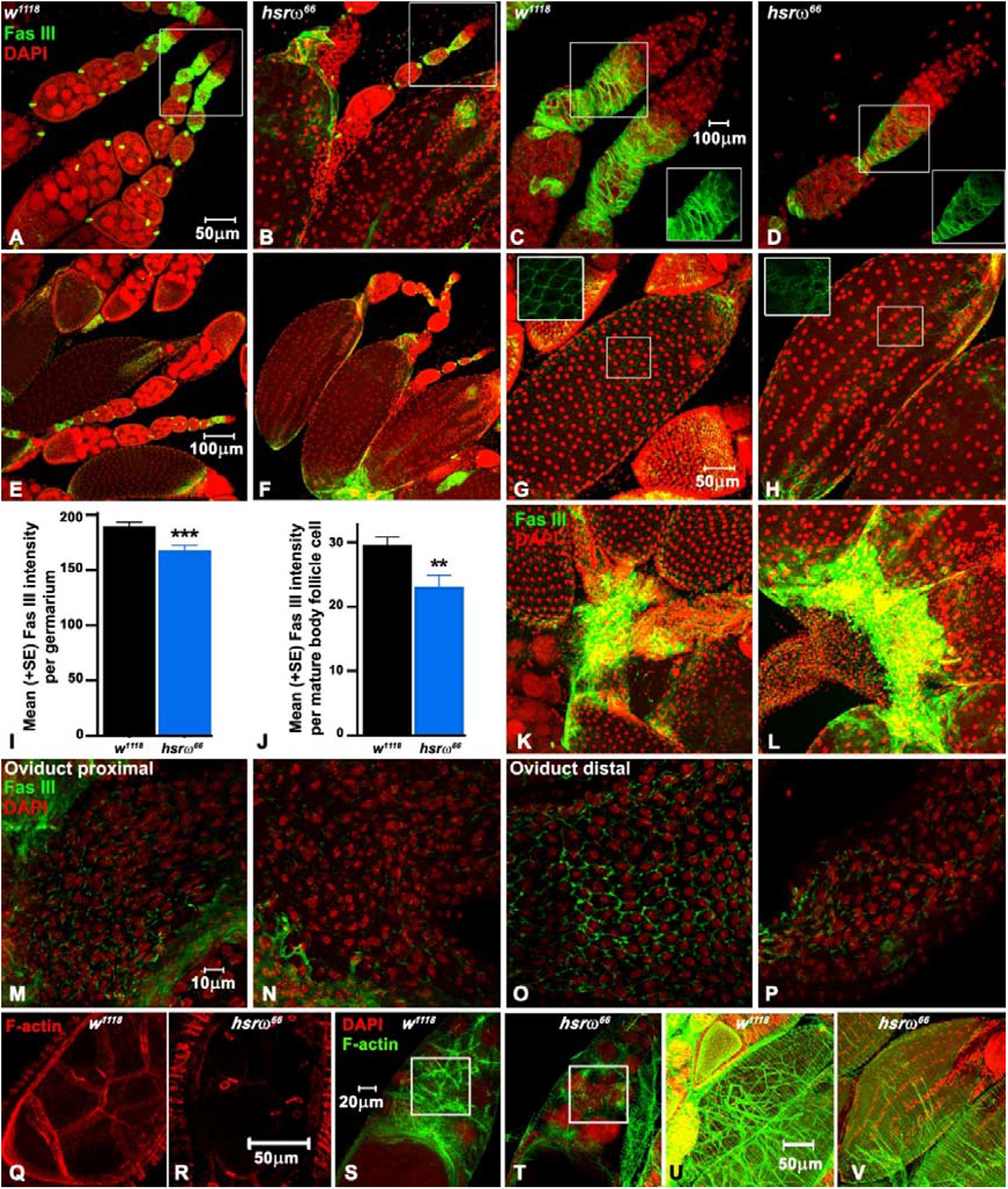
Fas III and F-actin levels are reduced in *hsr*ω*^66^* egg chambers. **(A-H)** Fas III distribution in germarium/early (**A-D**), mid (**E-F**) and late/mature stages (**G-H**) in 6 days old *w^1118^* (**A, C, E, G**) and *hsr*ω*^66^* (**B, D, F, H**); **C** and **D** are higher magnifications of the boxed regions in **A** and **B** respectively, with the insets (lower right) in each showing only Fas III staining of the boxed regions in respective panel; insets (upper left) in **G** and **H** are higher magnifications showing only Fas III staining in the boxed regions to reveal follicle cell arrangements in the two genotypes. **(I-J)** Fas III fluorescence intensities (arbitrary units, Y-axis) in germarium (**I**, N= 24 and 25 for *w^1118^* and *hsr*ω*^66^*, respectively) and mature body follicle cells (**J**, N= 7 for each genotype). (**K**-**L**) Fas III positive (green) corpus luteum at the ends of the mature follicles in *w^1118^* (**K**) and *hsr*ω*^66^* **(L)**; (**M P**) proximal (**M**, **N**) and distal (**O**, **P**) parts of the Fas III stained (green) lateral oviduct in *w^1118^* (**M**, **O**) and *hsr*ω*^66^* (**N**, **P**). **A-H** and **K-P** also show DAPI in red. (**Q**-**V**) Phalloidin stained F-actin (red in **Q, R** and green in **S-V**) distribution in stage 8 (**Q, R**), 10 (**S, T**) and 13/14 (**U, V**) egg chambers in *w^1118^* (**Q**, **S** and **U**) and *hsr*ω*^66^* (**R**, **T** and **V**); DAPI staining (red) is also shown in **S-V**. Scale bar in **A** (50µm) applies to **A-B,** in **C** (10µm) to **C-D**, in **E** (100µm) to **E-F**, in **G** (50µm) to **G-H** and **K**-**L,** in **M** (10µm) to **M-P**, in **R** (50µm) to **Q-R**, in **S** (20µm) to **S-T** while that in **U** (50µm) to **U-V**.

### Oogenesis defects in *hsr*ω*^66^* paralleled by targeted perturbations of *hsr*ω transcript levels

To confirm that the ovarian defects in *hsr*ω*^66^* are indeed due to compromised levels of various *hsr*ω lncRNAs, we examined effects of targeted down- or up-regulation of *hsr*ω transcripts in otherwise wild type background. For down-regulation, we used *hsr*ω*RNAi* transgene that targets the tandem-repeat containing *hsr*ω*RB*, *hsr*ω*RG*, and *hsr*ω*RF* transcripts (Mallik and Lakhotia, 2009) and *hsr*ω*exRNAi* (Sahu et al., 2021) transgene targeting exon 2 in the *hsr*ω lncRNAs, while for upregulation of these transcripts, the *EP93D* allele was used (Sengupta and Lakhotia, 2006). Additionally, the *hsr*ω*RH* transcript (www.flybase.org) was also expressed in *hsr*ω*^66^* background by recombining the *UAS hsr*ω*RH* transgene (Sahu et al., 2021) and *hsr*ω*^66^* allele. A follicle cell specific *tjGAL4* (Weaver et al., 2020), the ubiquitous *daGAL4* (Chaturvedi et al., 2016) and *ActGAL4* (Hongay and Orr-Weaver, 2011) drivers were used singly or in combination to drive these transgenes for examination of ovarian phenotypes in 6 days old females. The expected expression domains of the GAL4 drivers were confirmed through *UAS-EGFP* expression in pilot studies.

Ovaries from the *tjGAL4* (Fig. 3A), *daGAL4* (Fig. 3I) or *ActGAL4* (Fig. 3P) driver lines and each of the undriven transgene lines (Fig. 3B, D, G) showed normal oogenesis but down regulation of *hsr*ω transcripts resulted in manifestation of some or all the defects seen in *hsr*ω*^66^* ovaries. The *tjGAL4>hsr*ω*RNAi* (Fig. 3C), *tjGAL4>hsr*ω*exRNAi* (Fig. 3E) and *tjGAL4>hsr*ω*RNAi hsr*ω*exRNAi* (Fig. 3F) ovaries showed paucity of mid-stage chambers and ovulation block, which was greater when the *hsr*ω*exRNAi* was expressed alone or together with *hsr*ω*RNAi* (Fig. 3S). Interestingly, while the *EP93D* ovaries showed near normal ovaries (Fig. 3G), over-expression of *hsr*ω transcripts in *tjGAL4>EP93D* ovaries (Fig. 3H) also caused ovulation block and paucity of mid-stage chambers. Co-targeting the *EP93D* allele with *tjGAL4* and *daGAL4* ovaries (Fig. 3J) caused more severe ovulation block (also see Fig. 3S). For a summary of ovarian phenotypes in these genotypes, also see Table 1 (rows 1-7).

**Table 1.** Summary of ovary phenotypes in 6 days old females of different genotypes.

| Genotype | Ovary size & egg laying | Mid- stage chambers | Late-stage chambers | Ovulation block* | Modulation of ovarian phenotype |
| --- | --- | --- | --- | --- | --- |
| 1. *hsrω<sup>66</sup>* (Figs. 1,2,3) | Small, fewer eggs laid | Frequent apoptosis | Fewer | Moderate | Oogenesis disrupted |
| 2. *tjGAL4>hsrωRNAi* (Fig. 4C) | Normal | Fewer | Fewer | Mild | Mild disruption |
| 3. *tjGAL4>hsrωexRNAi* (Fig. 4E) | Normal, fewer eggs laid | Fewer | Fewer | High | Moderate disruption |
| 4. *tjGAL4>hsrωRNAi/hsrωexRNAi* (Fig. 4F) |  |  |  |  |  |
| 5. *tjGAL4>EP93D* (Fig. 4H) | Asymmetric ovaries, fewer eggs laid | Frequent apoptosis | Fewer | High | Moderate disruption |
| 6. *tjGAL4; daGAL4>EP93D* (Fig. 4J) |  |  |  |  |  |
| 7. *daGAL4>hsrωRNAi* (Fig. 4M) | Normal, fewer eggs laid | Apoptosis frequency high | Fewer | Mild | Mild disruption |
| **UAS-*hsrωRH* expression in *hsrω<sup>66</sup>* ovary** |  |  |  |  |  |
| 8. *tjGAL4>UAS-*hsrωRH* hsrω<sup>66</sup>* (Fig. 4O, V) | Small, fewer eggs laid | Fewer | Better than in *hsrω<sup>66</sup>* | Moderate | Mild rescue of *hsrω<sup>66</sup>* phenotypes |
| 9. *ActGAL4>UAS-*hsrωRH* hsrω<sup>66</sup>* (Fig. 4Q, W) | Normal, more eggs laid than *hsrω<sup>66</sup>* | Normal | Near normal | Mild (partial rescue) | Moderate rescue of *hsrω<sup>66</sup>* phenotypes |
| 10. *ActGAL4/tjGAL4>UAS-*hsrωRH* hsrω<sup>66</sup>* (Fig. 4R, X) | Normal, eggs laid as in wild type | Normal | Near normal | No | Substantial rescue of *hsrω<sup>66</sup>* phenotypes |
| **Interaction of N<sup>ACT</sup> with *hsrωRNAi* and *EP93D*** |  |  |  |  |  |
| 11. *tjGAL4>UAS-N<sup>act</sup>* (Fig. 6B) | Small, No eggs laid | Normal | No chambers >stage 9 | No mature follicles | Mid-oogenesis onwards disrupted |
| 12. *tjGAL4>UAS-N<sup>act</sup>/hsrωRNAi* (Fig. 6C) | Small, No eggs laid | Normal | No chambers >stage 9 | No mature follicles | No change following *hsrω* down- or up-regulation |
| 13. *tjGAL4>UAS-N<sup>act</sup>/EP93D* (Fig. 6D) |  |  |  |  |  |
| **Interaction of Cut with *hsrωRNAi* and *EP93D*** |  |  |  |  |  |
| 14. *tjGAL4>cutRNAi* (Fig. 6F) | Normal, very few small, rounded eggs laid | Antero-posterior axis reduced | Length reduced | High | No change by *hsrω* down-regulation |
| 15. *tjGAL4>cutRNAi/hsrωRNAi* (Fig. 6G) |  |  |  |  |  |
| 16. *tjGAL4>cutRNAi/EP93D* (Fig. 6H) | Normal, eggs laid | Near wild type | Length near wild type | Mild | Partial rescue |
| 17. *tjGAL4>UAS-cut* (Fig. 6J) | Small, no eggs laid | Germarium and early-stage chambers not identifiable | Abnormal vitellogenic chambers | No mature follicles | Oogenesis completely disrupted |
| 18. *tjGAL4>UAS-cut/hsrωRNAi* (Fig. 6K) | Small, no eggs laid | Germarium and early-stage chambers not identifiable | Abnormal vitellogenic chambers | No mature follicles | No change following *hsrω* down-regulation |
| 19. *tjGAL4>UAS-cut/EP93D* (Fig. 6L) | Larger, no eggs laid | Rescued to near wild | Abnormal chambers | No mature follicles | Rescue of early but not |
|  |  | type |  |  | mid- and late stages |
| **Interaction of TBPH with *hsrω*RNAi and *EP93D*** |  |  |  |  |  |
| 20. *tjGAL4>TBPHRNAi* (Fig. 7B) | Normal, fewer eggs laid | Normal | Normal | Moderate | NA |
| 21. *tjGAL4> TBPHRNAi/hsrωRNAi* (Fig. 7C) | Normal, eggs laid | Normal | Normal | Mild | Partial rescue |
| 22. *tjGAL4>TBPHRNAi/EP93D* (Fig. 7D) | Normal, very few eggs laid | Fewer | Fewer | High | Exaggerated |
| 23. *tjGAL4>UAS-TBPHGFP* (Fig. 7F) | Normal, no eggs laid | Normal | Late stage chambers do not mature | No mature follicles | No change following *hsrω* up-regulation |
| 24. *tjGAL4>TBPHGFP/EP93D* (Fig. 7H) |  |  |  |  |  |
| 25. *tjGAL4>TBPHGFP/hsrωRNAi* (Fig. 7G) | Normal, fewer eggs laid | Normal | Normal | Moderate | Moderate rescue after *hsrω* down-regulation |
| **Interaction of Caz with *hsrω*RNAi and *EP93D*** |  |  |  |  |  |
| 26. *tjGAL4>cazRNAi* (Fig. 7J) | Normal, smaller eggs laid | Normal | Length reduced | Normal ovulation | No change following *hsrω*-down regulation |
| 27. *tjGAL4>cazRNAi/hsrωRNAi* (Fig. 7K) | Normal, near normal size eggs laid | Normal | Length partially restored | Normal ovulation | Partial rescue following *hsrω* down- or up-regulation |
| 28. *tjGAL4>cazRNAi/EP93D* (Fig. 7L) |  |  |  |  |  |
| 29. *tjGAL4>UAS-FUS<sup>WT</sup>* (Fig. 7N) | Normal, no eggs laid | Normal | Fail to mature | No mature follicles | No change following *hsrω* up-regulation |
| 30. *tjGAL4>UAS-FUS<sup>WT</sup>/EP93D* (Fig. 7P) |  |  |  |  |  |
| 31. *tjGAL4>UAS-FUS<sup>WT</sup>/hsrωRNAi* (Fig. 7O) | Normal, eggs laid | Normal | Normal | Moderate | Rescue following *hsrω* down-regulation of |
| **Ecdysone feeding of *hsrω<sup>66</sup>* flies** |  |  |  |  |  |
| 32. 20HE fed *hsrω<sup>66</sup>* (8 days old) (Fig. 8E) | Ovary size and fecundity better in 80% cases | As in *hsrω<sup>66</sup>* | Better than *hsrω<sup>66</sup>* | Rescued in 60% cases | Substantial rescue |

\*Degree of ovulation block defined by average number of mature follicles/ovariole: Mild = 2; Moderate = 3; High = >3

The mid-oogenesis stage 8 is a check point for quality control of the developing egg chambers (Drummond-Barbosa and Spradling, 2001; Lebo and McCall, 2021). Unlike typical DAPI stained healthy nurse and follicle cell nuclei in *w^1118^* egg chambers of different stages (Fig. 3K), the *hsr*ω*^66^* ovaries frequently displayed fragmented nuclei in stage 6-8 nurse and/or follicle cells (Fig. 3L, white arrow), a characteristic feature of apoptosis. The *daGAL>hsr*ω*RNAi*, although not showing major ovarian defects, exhibited more frequent apoptotic mid-stage chambers (Fig. 3M), which was confirmed by quantification of apoptotic egg chamber frequency per ovariole in *w^1118^*, *hsr*ω*^66^* and *daGAL>hsr*ω*RNAi* ovaries (Fig. 3N).

We next checked if the defective oogenesis in *hsr*ω*^66^* ovaries can be rescued by supplementing some of the *hsr*ω transcripts through the inducible *UAS-hsr*ω*RH* transgene which expresses the RH transcript of *hsr*ω gene under *UAS* promoter (Sahu et al., 2021). Recombination of this chromosome 3 inserted transgene with the *hsr*ω*^66^* allele generated *UAS-hsr*ω*RH hsr*ω*^66^* stock. The *tjGAL4* or *ActGAL4* driven expression in *UAS-hsr*ω*RH hsr*ω*^66^* flies, led to some improvement in survival of early and mid-stage chambers, but the ovulation block persisted (Fig. 3O, S); the rescue was slightly more pronounced in *ActGAL4>UAS-hsr*ω*RH hsr*ω*^66^* (Fig. 3Q, S). Remarkably, driving *UAS-hsr*ω*RH* by a combination of *ActGAL4* and *tjGAL4* drivers in *ActGAL4/tjGAL4>UAS-hsr*ω*RH hsr*ω*^66^* females elicited substantial recovery in oogenesis (Fig. 3R), with the mean number of ovarioles in 6 days old *ActGAL4/tjGAL4>UAS-hsr*ω*RH hsr*ω*^66^* ovaries being 15.35 (±0.15 S.E.) and the mean % apoptotic egg chambers per ovariole 4.89 (±0.37 S.E.) as in *w^1118^* females. The ovulation block seen in *hsr*ω*^66^* homozygous ovaries was also nearly completely absent in these ovaries (Fig. 3S, also see Table 1, rows 8-10).

We examined Caz and F-actin distribution in *w^1118^*, *hsr*ω*^66^*, *tjGAL4>hsr*ω*RH hsr*ω*^66^*, *ActGAL4>hsr*ω*RH hsr*ω*^66^* and *ActGAL4/tjGAL4>UAS-hsr*ω*RH hsr*ω*^66^* ovaries (Fig. 3T-X’). Since the distribution of Caz protein, the fly homolog of human Fus (Singh, 2023), has not been described earlier in *Drosophila* ovaries, a brief description of its expression patterns in *w^1118^* and *hsr*ω*^66^* ovaries is first provided. Caz is expressed from germarium to mature follicles in *w^1118^* ovaries, (Fig. 3T-X’, Supplementary Fig. S2). Caz was present in germline as well as somatic cells till early egg chambers, (Fig. 3T, Supplementary Fig. S2) but during mid-oogenesis Caz appeared more abundant in follicle cells than in germline nurse cells (Fig. 3T, Supplementary Fig. S2). The ovarian sheath cells were strongly stained for Caz. In stage 14 mature *w^1118^* follicles, the posterior follicle cells, which undergo trimming for ovulation follicle, expressed Caz at higher level than the body follicle cells (Supplementary Fig. S2B). Corpus luteum of mature follicles also showed high levels (Fig. Supplementary Fig. S2B). As in other tissues, Caz was always nuclear in ovarian cells. Caz expression in *hsr*ω*^66^* germarium (Fig. 3U and Supplementary Fig. S2C, C’) was generally comparable with *w^1118^* although the staining intensity appeared weaker in all cell types. Unlike the forward movement and consequent sparse distribution of anterior follicle cells in early stage 14 follicles in *w^1118^* ovarioles, these cells in similar stage *hsr*ω*^66^* chambers remain aggregated at base of the anterior end (insets in Supplementary Fig. S2B, D) although with comparable Caz staining. Caz expression in anterior follicle cells of late stage 14 was weaker in *hsr*ω*^66^* than in *w^1118^* (Supplementary Fig. S2B, D).

In agreement with the above noted ovarian phenotypes, Caz expression in *tjGAL4>hsr*ω*RH hsr*ω*^66^* ovaries displayed only a marginal improvement (Fig. 3V, V’), but in *ActGAL4>hsr*ω*RH hsr*ω*^66^* ovaries, Caz and F-actin levels and distribution of follicle cell nuclei in (Fig. 3W, W’) appeared substantially better. In agreement with the near normal ovarian organization in *ActGAL4/tjGAL4>hsr*ω*RH hsr*ω*^66^* ovaries (Fig. 3R), the distribution of Caz, F-actin and follicle cell nuclei (Fig. 3X, X’) appeared remarkably similar to that in *w^1118^* ovaries.

Taken together, these results confirm that the diverse oogenesis defects seen in *hsr*ω*^66^* are indeed related to the perturbed levels of this gene’s lncRNAs.

**Fig. 3.**
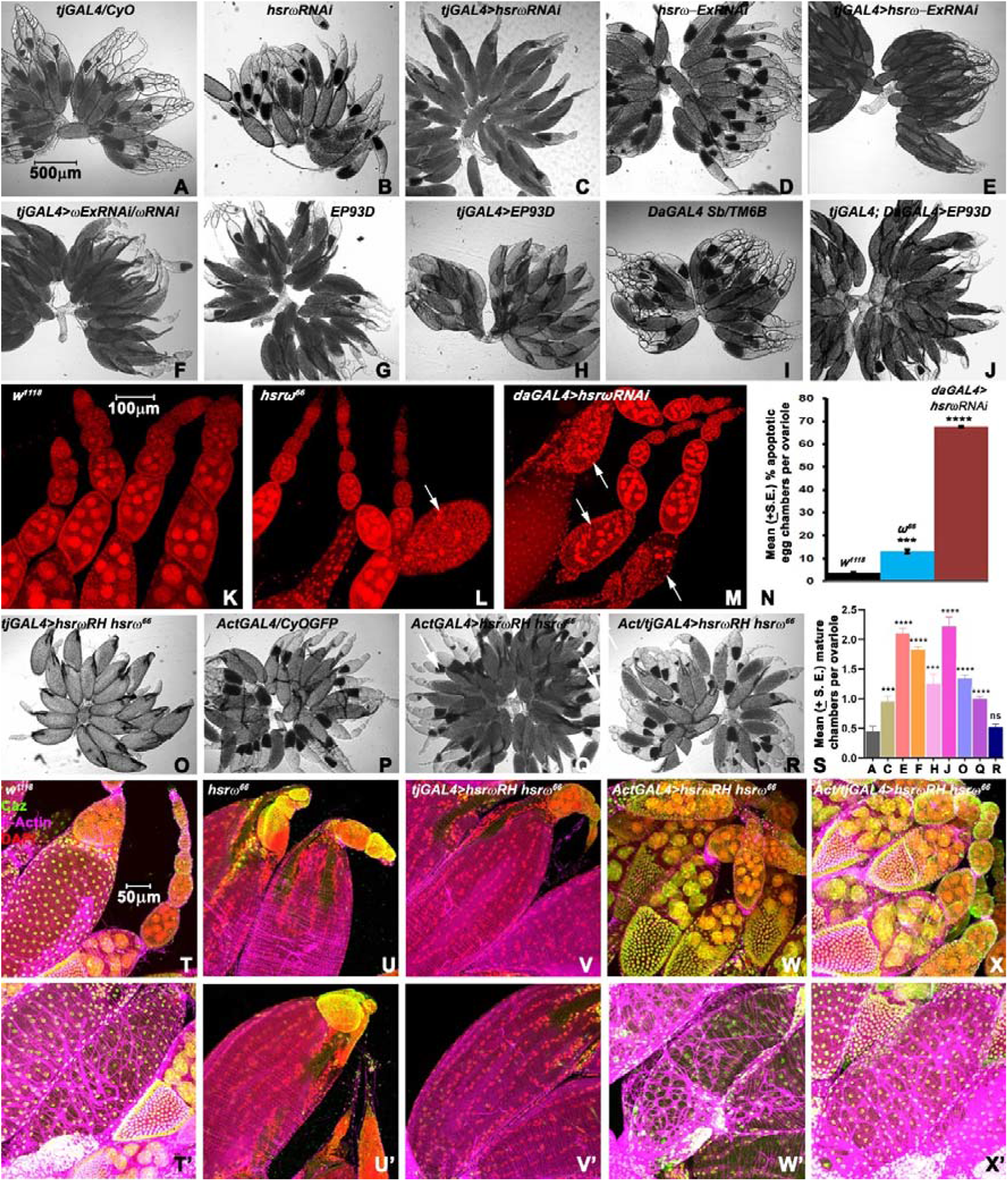
Defects in oogenesis in *hsr*ω*^66^* ovaries are due to perturbations in levels of *hsr*ω transcripts. (**A**-**J**) Ovaries of different genotypes noted on top in each panel. (**K-M**) DAPI (red) stained ovarioles of different genotypes (noted on top of each panel); white arrows indicate apoptotic chambers. (**N**) Mean % apoptotic egg chambers per ovariole (Y-axis; N = 60 ovaries for each genotype)) in genotypes noted above each bar. (**O**-**R**) Ovaries from 6 days old females of different genotypes noted on top in each panel. (**S**) Mean numbers of mature follicles per ovariole (Y-axis, N=18 ovaries for **A** and 20 ovaries for all the other genotypes) in different genotypes (X-axis) indicated by the corresponding panel number in this figure. (**T**-**X’**) Distribution of Caz (green) and F-actin (magenta) in early and mid (**T**-**X**) and late (**T’**-**T’**) stage egg chambers s of different genotypes (noted on top of each column); DAPI staining is in red. Scale bar in **A** (500µm) applies to **A-J** and **O R**, in **K** (500µm) to **K-M** and that in **T** (50µm) to **T-X’**.

### Cut, but not Hindsight, expression reduced during late oogenesis in *hsr*ω*^66^* ovarioles

Since Cut and Hindsight (Hnt) play important roles in the ovulation process (Knapp et al., 2019), we compared distribution of these transcription factors in *w^1118^* and *hsr*ω*^66^* late stages egg chambers (Fig. 4).

Cut is expressed transiently in main body follicle cells during stages 10B-13 but disappears in stage 14 mature follicles (Knapp et al., 2019). Compared to the *w^1118^* (Fig. 4F, H, and J), levels of Cut were low even in stage 10-13 *hsr*ω*^66^* chambers (Fig. 4A-F). During these stages, the highest level of Cut was observed in the centripetal follicle cells of *w^1118^* chambers, with the main body follicle cells being also Cut-positive (Fig. 4C, E); however, immunostaining for Cut in *hsr*ω*^66^* stage 10-13 egg chambers was significantly lower in centripetal follicle cells at all these stages, and nearly absent in body follicle cells between stages 10 and 13 (Fig. 4B, D, F).

Hnt is not expressed in stage 11-13 follicle cells but its presence in the mature stage 14 follicles is essential for follicle rupture and ovulation competency (Knapp et al., 2019). Compared to follicle cells of 7-10A stage *w^1118^* chambers (Fig. 4G), the Hnt staining was relatively weak in the few *hsr*ω*^66^* mid-stage chambers (Fig. 4H) present in these ovaries. The stage 14 mature follicles in *w^1118^* and *hsr*ω*^66^* ovaries, however, showed more or less comparable Hnt levels, with higher levels in anterior (Fig. 4I, J) and posterior follicle cells (Fig. 4K, L) in both.

**Fig. 4.**
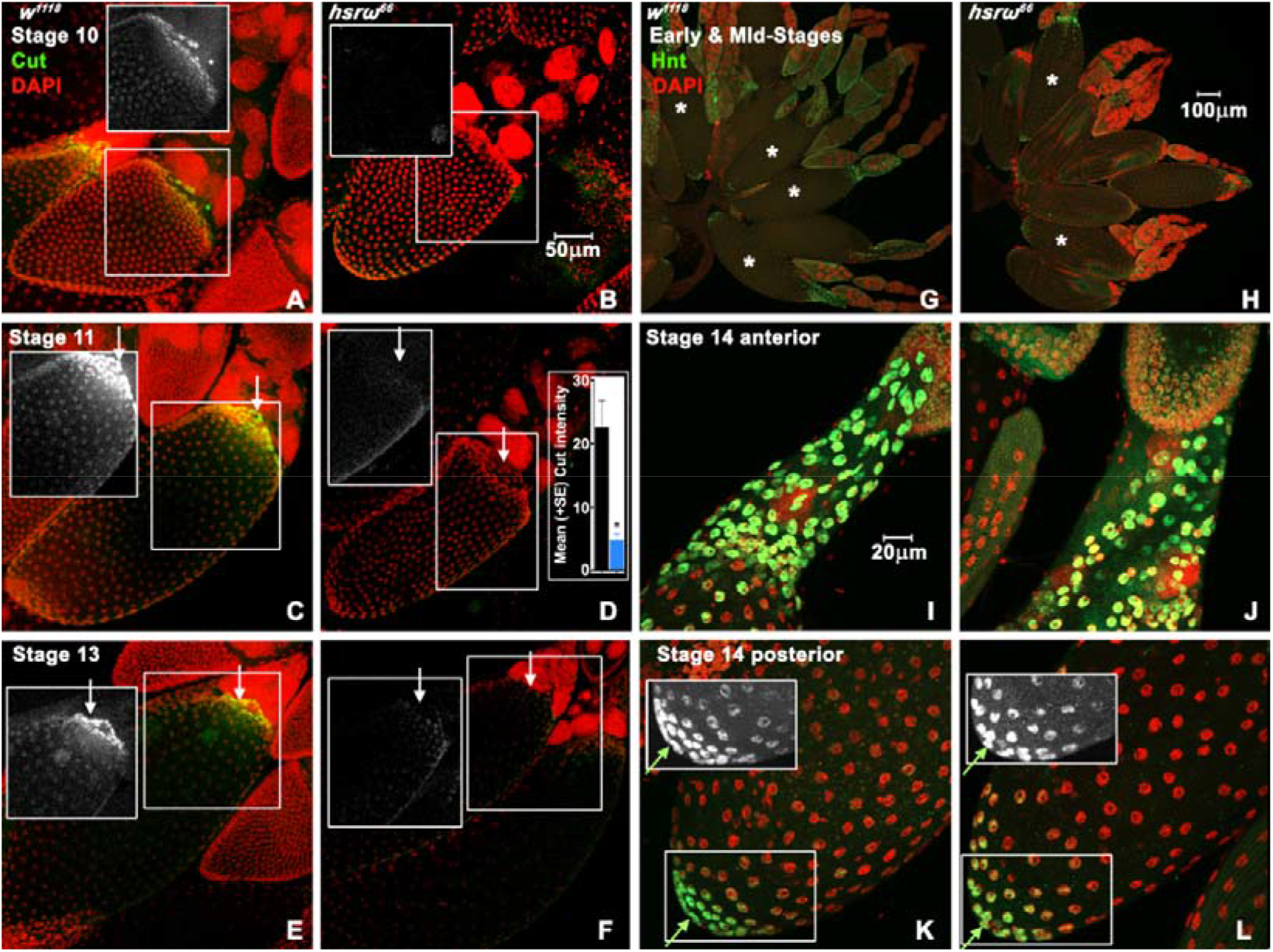
Near absence of *hsr*ω transcripts reduces Cut but not Hindsight (Hnt) during late oogenesis. **(A-F)** Distribution of Cut (green) in stage 10 (**A, B**), 11 (**C, D**) and 13 (**E**, **F**) follicles from *w^1118^* (**A, C, E**) and *hsr*ω*^66^* ovarioles (**B, D, F**); DAPI in red; insets show only the Cut fluorescence (white) in respective boxed areas with white arrows indicating centripetal follicle cells; bar graph in mid-right inset in **D** shows Cut fluorescence intensity (in arbitrary units, Y-axis) per centripetal follicle cell in *w^1118^* (black bar) and *hsr*ω*^66^* (blue bar) stage 11 egg chamber (N= 70 and 50 cells from 7 and 5 *w^1118^* and *hsr*ω*^66^* chambers, respectively). **(G-L)** Hnt (green) and DAPI (red) staining in early and mid-stage (**G**, **H**) and in anterior (**I-J**) and posterior (**K-L**) parts of stage 14 egg chambers from *w^1118^* (**G, I, K**) and *hsr*ω*^66^* **(H, J, L)**; insets in **K**, and **L** show only Hnt (white) staining in the boxed areas in the respective panels while white arrows indicate the Hnt positive posterior follicle cells. Scale bar in **B** (50µm) applies to **A**-**L**.

### Potential genetic interactors of *hsr*ω during oogenesis

With a view to identify possible interactors and/or targets of the *hsr*ω lncRNAs, we selected two candidates, the Notch and Cut, which are major regulators of *Drosophila* oogenesis and ovulation (Deady et al., 2017; Jackson and Blochlinger, 1997; Sun and Deng, 2005) and two hnRNPs, the TBPH/TDP-43 and Caz/dFus which are known to physically interact with the *hsr*ω nuclear RNA (Singh, 2023) for genetic interaction studies. We examined effects of *tj GAL4* driven altered levels of these proteins in conjunction of down- or up-regulation of *hsr*ω transcripts. For summary of ovarian phenotypes in these genotypes, see Table 1 (rows 11-31).

The undriven *UAS-N^Act^* did not affect oogenesis (Fig. 5A). The *tjGAL4>N^Act^* ovaries showed resulted in complete absence of egg chambers beyond stage 9 (Fig. 5B). Down- or up regulation of *hsr*ω transcripts did not modify the *tjGAL4>N^Act^* phenotype since *tjGAL4>N^Act^/hsr*ω*RNAi* or *tjGAL4>UAS-N^Act^/EP93D* ovaries did not show any egg chamber beyond stage 9 (Fig. 5C, D). As expected, none of these females laid any eggs (Table 1, rows 11-13). Absence of any modulation of the severely disrupted oogenesis in *tj-GAL4>N^Act^* ovaries by down- or up-regulation of *hsr*ω transcripts suggests that Notch signaling is upstream of action/s of these transcripts.

The *cutRNAi* transgene by itself did not affect oogenesis (Fig. 5E, E’). Expression of *tjGAL4*>*cutRNAi* in wild type *hsr*ω background did not affect ovary size or ovariole numbers (Fig. 5F) and early egg chambers. However, the antero-posterior (A-P) axis of egg chambers was significantly reduced from mid-stage onwards resulting in smaller and rounded mature follicles with aberrant dorsal appendages (insets in Fig. 5F); these ovaries also displayed significant apoptosis in mid-stage egg chambers (Fig. 5F’), disrupted follicle cell arrangement (inset in Fig. 5F’) and ovulation block (Fig. 5F). Ovarioles suffering co-down regulation of Cut and *hsr*ω transcripts (*tjGAL4*>*cutRNAi/hsr*ω*RNAi*) appeared generally like *tjGAL4*>*cutRNAi*. The follicle cell layer covering the mature roundish follicles was disorganized in both these genotypes (insets in Fig. 5F-G). Surprisingly, in both cases, females laid roundish eggs which developed into viable larvae (data not presented). Interestingly, ovaries in *tjGAL4*>*cutRNAi/EP93D* appeared significantly better since mid stage chambers were more frequent than in the previous three genotypes and the organization of follicle cell layer and length of mature follicles and of laid eggs were nearly like those in wild type; the ovulation block was also substantially rescued (Fig. 5H, H’, also see Table 1, rows 14-16).

The *UAS-cut* transgene by itself did not affect oogenesis (Fig. 5I, I’) but *tjGAL4*>*UAS-cut* ovaries showed gross disorganization of ovarioles as germarium and developing egg chambers lost their characteristic organization, with aberrant location of yolk in vitellogenic stages, so that no mature follicles formed (Fig. 6J, J’). The *tjGAL4*>*cut; hsr*ω*RNAi* ovarioles were also disorganized (Fig. 5K, K’) as in *tjGAL4*>*cut* ovaries. Surprisingly, *tjGAL4* driven co-over-expression of Cut and *hsr*ω transcripts (*tjGAL4*>*cut; EP93D*) significantly improved the ovarian organization till the mid-stage egg chambers although many late-stage chambers showed apoptosis (Fig. 5L, L’) resulting in complete absence of mature follicles. No eggs were laid by females with *tjGAL4* driven up-regulated Cut expression, without or with altered levels of *hsr*ω transcripts (Table 1, rows 17-19).

**Fig. 5.**
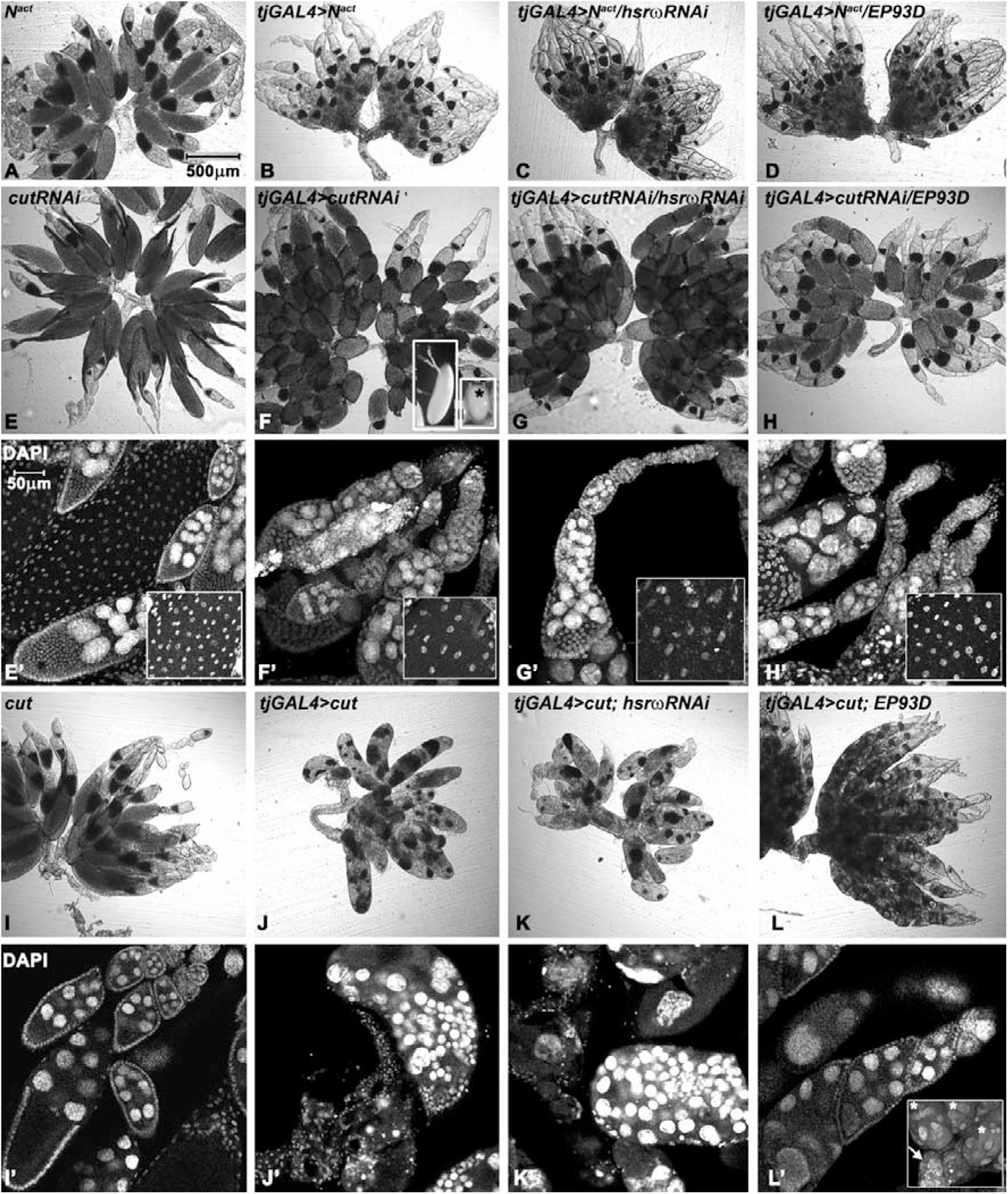
The *tjGAL4* driven mis-expression of active Notch in follicle cells disrupts later stages of oogenesis irrespective of levels of *hsr*ω transcripts while over-expression of *hsr*ω partially rescues effects of down or up-regulation of Cut in follicle cells. **(A-D)** DIC images of undriven *UAS-N^Act^* (**A**), *tjGAL4>N^Act^* (**B**), *tjGAL4> N^Act^/hsr*ω*RNAi*, (**C**), and *tjGAL4>N^Act^/EP93D* (**D**) ovaries. **(E-H’)** DIC (**E-H**) and DAPI-stained (white) (**E’-H’**) confocal images of undriven *UAS-cutRNAi* (**E, E’**), *tjGAL4>cutRNAi* (**F, F’**), *tjGAL4>cutRNAi/hsr*ω*RNAi*, (**G, G’**), and *tjGAL4>cutRNAi/EP93D* (**H, H’**) ovaries; insets in **E’-H’** show magnified images of body follicle cells in late-stage chambers. (**I-L’**) DIC (**I-L**) and DAPI-stained (white) confocal optical sections (**I’-L’**) of undriven *UAS-cut* (**I, I’**), *tjGAL4>cut* (**J, J’**), *tjGAL4>cut/hsr*ω*RNAi*, (**K, K’**), and *tjGAL4>cut/EP93D* (**L, L’**) ovaries; inset in **L’** shows DAPI (white) stained projection image of junction of ovarioles and lateral oviduct (white arrow) with abnormal late stage follicles (* marked). Scale bar in **A** (500µm) applies to **A-H** and **I-L** and that in **E’** (50µm) to **E’-H’** and **I’-L’**.

Undriven *TBPHRNAi* transgene did not affect oogenesis (Fig. 6A) but it’s down-regulation with *tjGAL4* (*tjGAL4>TBPHRNAi*) caused ovulation block with insignificant effect on early and mid-stage chambers (Fig. 6B). Interestingly, co-down regulation of TBPH and *hsr*ω transcripts (*tjGAL4>TBPHRNAi; hsr*ω*RNAi*) rescued ovulation block (Fig. 6C) so that these flies laid normal eggs. Upregulation of *hsr*ω in *TBPHRNAi* background (*tjGAL4>TBPHRNAi; hsr*ω*RNAi*) enhanced ovulation block (Fig. 6D, also see Table 1, rows 20-22).

Oogenesis in the *UAS-TBPHGFP* transgene carrying females was normal (Fig. 6E) but *tjGAL4>TBPHGFP* lacked mature late-stage chambers although earlier stages looked normal (Fig. 6F). Co-expression of *hsr*ω*RNAi* in this background (*tjGAL4>TBPHGFP hsr*ω*RNAi*) promoted follicle maturation but they suffered moderate ovulation block (Fig. 6G). Interestingly, the *tjGAL4>TBPHGFP EP93D* ovaries were similar to *tjGAL4>TBPHGFP* ovaries and lacked mature follicles (Fig. 6H, also see Table 1 rows 23-25).

The *cazRNAi* transgene by itself did not affect oogenesis (Fig. 6I) but it’s down-regulation in *tjGAL4*>*cazRNAi* ovaries caused reduction in mature follicle length (Fig. 6J) although these eggs were laid and hatched. Interestingly, *tjGAL4*>*cazRNAi* ovaries with down- or up regulated *hsr*ω transcripts (*tjGAL4*>*cazRNAi*/*hsr*ω*RNAi* or *tjGAL4*>*/EP93D*, respectively) appeared better since lengths of mature follicles and eggs were closer to wild type (Fig. 6L, also see Table 1 rows 26-28).

Undriven wild type human *UAS-Fus^WT^* carrying stock displayed normal ovaries (Fig. 6M) but the transgene’s over-expression in *tjGAL4>Fus^WT^* ovaries prevented maturation of late follicles (Fig. 6N). Interestingly, co-expression of *hsr*ω*RNAi* (*tjGAL4>Fus^WT^ hsr*ω*RNAi*) or *EP93D* (*tjGAL4>Fus^WT^ EP93D*) generated phenotypes like that in *tjGAL4>TBPHGFP hsr*ω*RNAi* and *tjGAL4>TBPHGFP EP93D*, respectively, with follicles maturing but remaining unovulated or late follicles not maturing, respectively (Fig. 6O, P also see Table 1 rows 29-31).

**Fig. 6.**
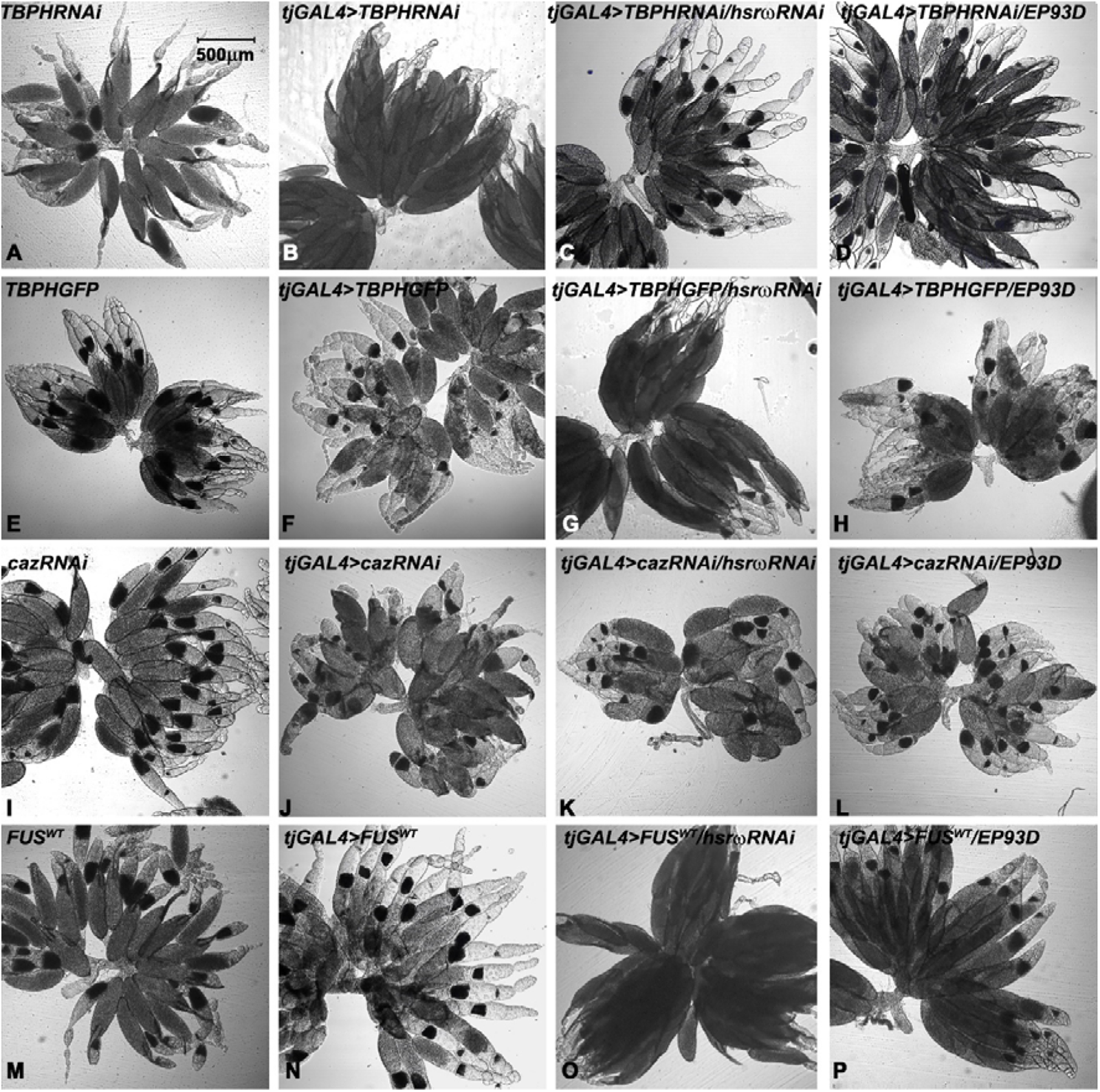
Varying effects of altered levels of *hsr*ω on ovarian phenotypes following *tjGAL4* driven down- or up-regulation of TBPH/TDP-43 or Caz/Fus in follicle cells. **(A-P)** DIC images of ovaries of different genotypes (noted on upper left corner in each panel). Scale bar in **A** (500µm) applies to **A-P**.

### Partial rescue of ovulation block in *hsr*ω*^66^* females by dietary 20-hyrdoxy-ecdysone (20-HE)

Since ecdysteroids are crucial for *Drosophila* oogenesis and ovulation (Knapp and Sun, 2017) and since the *hsr*ω transcripts are very abundant in ecdysone synthesizing prothoracic gland (Mutsuddi and Lakhotia, 1995), we examined levels of 20-HE in *w^1118^* and *hsr*ω*^66^* females. 20-Immunoassay for 20-HE in late female pupae and 6 days old *w^1118^* and *hsr*ω*^66^* flies revealed that while this hormone’s titers in late pupal stage were comparable in the two genotypes (Fig. 7A), the *hsr*ω*^66^* homozygous females showed significantly reduced 20-HE (Fig. 7A’). Interestingly, 20-HE levels in *hsr*ω*^66^*/*TM6B* heterozygous females, which show normal fertility and fecundity, were similar to that in *w^1118^* females (Fig. 7A’).

In view of the reduced levels of ecdysone *hsr*ω*^66^* homozygous females, the *w^1118^* and *hsr*ω*^66^* male and female flies were fed for 115 hours on 20-HE (0.05mM) supplemented food from 3^rd^ to 8^th^ day post-eclosion. Examination of ovaries on day 8 showed that dietary 20-HE had no adverse effect on oogenesis in *w^1118^* females as revealed by ovary morphology, DAPI and F-actin stating (Fig8B-D). Interestingly, however, the 20-HE fed *hsr*ω*^66^* females showed substantially improved progression of oogenesis in females since out of the 50 ovaries examined from 20-HE-fed *hsr*ω*^66^* females (N=25), 30 were without ovulation block while 40 showed normal looking mid-stage egg chambers (Fig. 7E-G, also see Table 1, row 32), which, as noted above, are rare in *hsr*ω*^66^*. The F-actin staining, however, remained weaker in all oogenesis stages in 20-HE fed flies (Fig. 7D and 8G). Compared to the normally grown *hsr*ω*^66^* females. the 20-HE fed flies survived better and laid more eggs (data not presented).

**Fig. 7.**
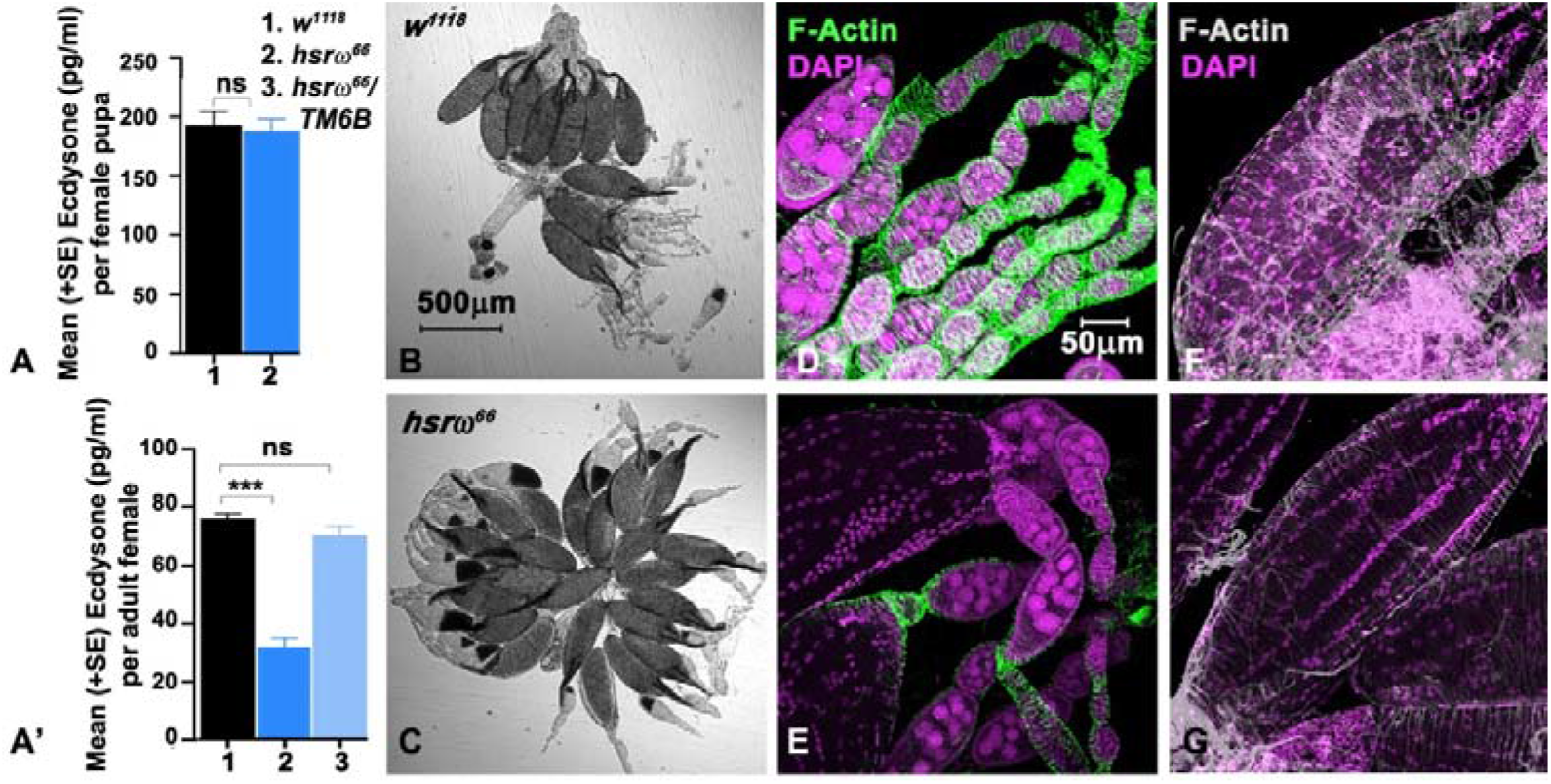
Reduced 20-HE levels in *hsr*ω*^66^* homozygous females and significantly improved oogenesis following dietary 20-HE. **(A-A’)** 20-HE titres (Y-axis) in female pupae (**A**) and 6 days old females (**A’**) of different genotypes (X-axis, the numbered genotypes are identified on top right corner). (**B-C**) DIC images of ovaries from 8 days old *w^1118^* (**B**) and *hsr*ω*^66^* (**C**) 20-HE fed females. (**D-G**) Phalloidin (green, F-actin) and DAPI (magenta) stained early- and mid-(**D-E**), and late-stage (**F-G**) egg chambers from 20HE fed *w^1118^* (**D, F**) and *hsr*ω*^66^* (**E, G**) females. Scale bar in **B** (500µm) applies to **B**-**C** and in **D** (50µm) to **D-G**.

## Discussion

Mis-expression of the multiple lncRNA encoding *hsr*ω gene of *Drosophila melanogaster* exerts pleiotropic effects on different life-history parameters and cell stress response (Lakhotia et al., 2012; Mallik and Lakhotia, 2011; Singh and Lakhotia, 2015). Present observation that life span and female fecundity are significantly reduced in *hsr*ω*^66^* homozygous flies that have little *hsr*ω transcripts (Ray et al., 2019a) agrees with earlier reports that loss of *hsr*ω causes high embryonic lethality and the few survivors are weak and short-lived (Johnson et al., 2011; Lo Piccolo and Yamaguchi, 2017; Mohler and Pardue, 1982; Ray and Lakhotia, 1998). Significantly, *hsr*ω is also identified as a major gene in normal aging because these transcripts greatly decline with age (Davie et al., 2018). Further reduced life span and fecundity of *hsr*ω*^66^* females when mated with *w^1118^* males is intriguing. Mating affects female physiology, including life span, through pheromones and seminal fluid proteins (Barnes et al., 2008; Koliada et al., 2020; Patlar and Civetta, 2022). Further studies are needed to see if *hsr*ω*^66^* females are sensitive to *w^1118^* male’s behaviour or seminal fluid components or both, although the normal fertility of *hsr*ω*^66^* males indicates non-essentiality of these transcripts for spermatogenesis.

As expected from high expression of *hsr*ω in most cell types of wild type ovary (Mutsuddi and Lakhotia, 1995), targeted down-regulation of *hsr*ω transcripts in ovaries recapitulated one or the other defects seen in *hsr*ω*^66^* ovaries. The stronger ovulation block in flies expressing *tjGAL4* driven *hsr*ω*exRNAi* singly or in combination with *hsr*ω*RNAi* compared to the milder defects following *tjGAL4* or *dalGAL4* driven *hsr*ω*RNAi* transgene expression correlates with their targets: while *hsr*ω*RNAi* majorly targets the 280bp repeat containing *hsr*ω nuclear transcripts (Mallik and Lakhotia, 2009), the *hsr*ω*exRNAi* targets exonic region, common to all *hsr*ω transcripts (Sahu et al., 2021). Commonality of oogenesis defects following reduced (*hsr*ω^66^, or *tjGAL4* or *dalGAL4* driven *hsr*ω*RNAi/hsr*ω*exRNAi*) as well as elevated (*tjGAL4>EP93D* or *tjGAL4; daGAL4>EP93D*) levels of these transcripts appear contradictory. However, since a major action of *hsr*ω transcripts is to regulate dynamics of diverse hnRNPs (Lakhotia, 2011; Lakhotia et al., 2012; Lo Piccolo et al., 2014; Prasanth et al., 2000; Singh and Lakhotia, 2015), disruptions of *hsr*ω gene activity are likely to have wide-ranging pleiotropic consequences which in some contexts can be, and are known to be similar irrespective of down-or up-regulation of these transcripts (Lakhotia, 2011; Mallik and Lakhotia, 2011; Ray and Lakhotia, 2019; Ray et al., 2019b). The substantially normal oogenesis following *tjGAL4* and *ActGAL4* driven expression of the *UAS-hsr*ω*RH* transgene in *hsr*ω*^66^* background firmly establishes that the near complete absence of *hsr*ω transcripts underlies the oogenesis defects in *hsr*ω*^66^* flies.

The delayed and poor egg-laying capacity of *hsr*ω*^66^* young females relates with small ovaries and fewer ovarioles in freshly emerged flies and slower follicle development. Apparently, the reduced *hsr*ω transcripts delay the initial phase of follicle development till 3 days post eclosion, while the ovulation block in older flies suppresses development of early/mid-stage follicles through feedback regulation (Drummond-Barbosa and Spradling, 2001; Meiselman et al., 2018). High apoptosis in mid-stage follicles associated with reduced *hsr*ω transcripts also adds to paucity of stage 7-13 chambers in >6 days old *hsr*ω*^66^* females. Besides being consequences of altered *hsr*ω transcript levels in ovary, the reduced ecdysone titer in *hsr*ω*^66^* females may also contribute to these phenotypes (König et al., 2011; Terashima et al., 2005).

A major action of the *hsr*ω nuclear lncRNAs is mediated through modulation of intra-cellular dynamics of RNA binding proteins, especially the hnRNPs (Lakhotia, 2011; Lakhotia et al., 2012; Lo Piccolo et al., 2014; Prasanth et al., 2000; Singh and Lakhotia, 2015). Since hnRNPs regulate gene expression, RNA processing and RNA transport (Chaudhury et al., 2010; Lin et al., 2015; Lo Piccolo et al., 2014) and since several hnRNPs have significant roles at specific stages of *Drosophila* oogenesis (Bansal et al., 2020; Casale et al., 2019; Dutta et al., 2020; Finger et al., 2022; Lasko, 2003; Peng and Gavis, 2022; Singh and Lakhotia, 2012), perturbed hnRNP dynamics following altered *hsr*ω transcript levels would cause wide pleiotropic consequences. Down-regulation of *hsr*ω transcripts affects sub cellular localization of TBPH/TDP-43 and Caz/dFus, the two recently identified hnRNP interactors of *hsr*ω and implicated in human syndromes with weak neuromuscular junctions (Chung et al., 2018; Lo Piccolo and Yamaguchi, 2017; Wang et al., 2011). The altered _metabolism of proteins like TBPH/TDP-43 and Caz/dFus in *hsr*ω_ *^66^* flies with greatly reduced *hsr*ω transcripts may also contribute to their weak neuro-muscular activities and lifespan. Indeed *hsr*ω*^66^* individuals display mobility defects and poor feeding-ability (data not shown), which also contribute to poor oogenesis since compromised nutrition and poor ovariole musculature affect oogenesis (Drummond-Barbosa and Spradling, 2001; Liao and Nässel, 2020; Meiselman et al., 2018).

Since fusomes regulate the pattern and synchrony of divisions during egg chamber differentiation, movement of mitochondria to oocyte, and several signaling events during follicle growth (Cox and Spradling, 2003; de Cuevas et al., 1996; Lighthouse et al., 2008; Lilly et al., 2000), weak fusomes and reduced Vasa in early-stage cysts in *hsr*ω*^66^* ovarioles would have multiple long-range consequences.

Reduced Cut in *hsr*ω*^66^* ovarioles may be due to lower 20-HE in these flies since ecdysone triggers the cascade that upregulates Cut after stage 10B (Knapp et al., 2019). Since Cut regulates soma-to-germline signaling, follicle cell migration and chorion gene amplification (Jackson and Blochlinger, 1997; Knapp et al., 2019), it’s reduction in *hsr*ω*^66^* may be responsible for disorganized arrangement of body follicle cells and their larger nuclei in late stage *hsr*ω*^66^* chambers. A mutual regulatory circuit between Cut and *hsr*ω transcripts is also evident from our genetic interaction results. Since targeted reduction in *hsr*ω transcript levels (*tjGAL4>hsr*ω*RNAi*) failed to modify the *tjGAL4>CutRNAi* as well as *tjGAL4>UAS-Cut* ovarian phenotypes, we believe that both low and high levels of Cut in follicle cells reduce *hsr*ω transcripts so that while their further down-regulation remains ineffective, their up regulation through *EP93D* co-expression partially rescues some of the *tjGAL4>CutRNAi* and *tjGAL4>UAS-Cut* ovarian phenotypes (Table 1, rows 14-20). Further studies would reveal if the Cut and *hsr*ω transcripts interact directly or through action/s of one or more of the *hsr*ω interacting hnRNPs.

Very little is known about roles of TBPH/TDP-43 and Caz/dFus in *Drosophila* oogenesis. Flybase (www.flybase.org) reports both these genes to express at low/moderate levels in adult ovary. A transcriptomic study also reported *caz* transcripts in ovary (Brown et al., 2014). Present study shows that Caz is indeed widely distributed in germinal as well as somatic cells of the germarium, and early-stage cysts, but is mostly limited to somatic follicle cells in mid- and late-stage wild type chambers. Reduced Caz in *hsr*ω*^66^* ovaries may reflect the reported degradation of Caz when *hsr*ω transcripts are low (Lo Piccolo et al., 2019).

The diverse defects in oogenesis following *tjGAL4* driven down- or up-regulation of TBPH or Caz in follicle cells (Table 1, rows 21-32) confirms that both these hnRNPs are required during oogenesis. Caz and TBPH are reported to function together with Caz being downstream (Wang et al., 2011). In agreement, the consequences of *tjGAL4* driven down- or up-regulation of TBPH or Caz in follicle cells with normal or down- or up-regulated *hsr*ω transcripts were mostly common. Different outcomes in some instances suggest that they may also act independently.

The diversity of consequences of modulation of Cut, TBPH or Caz levels in background of varying levels of *hsr*ω transcripts (Table 1) reflects complexity of interaction between the *hsr*ω transcripts and its multiple interacting proteins so that down or up-regulation of these transcripts may not always show the expected opposing effects, and in some cases, the two contrasting conditions may actually result in similar final outcome although through different paths (Bajpai et al., 2021; Lakhotia, 2011; Ray and Lakhotia, 2019; Ray et al., 2019b).

The poor general physiology of the surviving *hsr*ω*^66^* homozygotes can also affect progression of *Drosophila* oogenesis, which is highly dependent upon nutritional status and diverse paracrine and endocrine signaling (Drummond-Barbosa and Spradling, 2001; Kurogi et al., 2021; Liao and Nässel, 2020; Meiselman et al., 2018; Okamoto and Watanabe, 2022; Song and Zhou, 2020; Weaver and Drummond-Barbosa, 2021). The *hsr*ω gene is abundantly expressed in prothoracic gland (Mutsuddi and Lakhotia, 1995) and affects ecdysone as well Dilp8 signaling (Ray and Lakhotia, 2019; Ray et al., 2019b), both of which influence oogenesis and ovulation in *Drosophila* (Finger et al., 2021; König et al., 2011; Kurogi et al., 2021; Liao and Nässel, 2020; Meiselman et al., 2018; Terashima et al., 2005; Weaver and Drummond-Barbosa, 2021). Our present results show reduced levels of ecdysone in *hsr*ω*^66^* homozygotes. Therefore, the poor oogenesis in *hsr*ω*^66^* ovaries also reflects dysregulated intra- as well as inter-organ communications. The significant improvement in ovarian organization and oogenesis in majority of *hsr*ω*^66^* flies fed on ecdysone supplemented food confirms that reduced ecdysone metabolism does contribute to the various ovarian defects in flies with deficient *hsr*ω transcripts. Since exogenous ecdysone rescued most of the oogenesis defects in *hsr*ω*^66^* ovaries, the ecdysone receptors, whose mis-function can also affect oogenesis (Weaver and Drummond-Barbosa, 2021), are perhaps not affected by perturbed levels of *hsr*ω transcripts. Further studies are required to know if the ecdysone production in brain or ovaries or in both is affected in *hsr*ω mis-expressing individuals.

The *hsr*ω gene produces at least seven transcripts (www.flybase.org) using two transcription start and four termination sites and variable splicing of its single intron (Sahu et al., 2020). The *hsr*ω*^66^* homozygotes show extremely low levels of all these transcripts (Ray et al., 2019a). With a view to understand if all or some of these multiple transcripts have roles in *Drosophila* oogenesis, we examined, as a first step, effects of expression of the *UAS-hsr*ω*RH* transcripts in *hsr*ω*^66^* ovaries. The RH transcript corresponds to the region between the second transcription start site (TSS2) and second termination site located just upstream of the 280bp tandem repeats (Sahu et al., 2020). Since the *tjGAL4>hsr*ω*RNAi* or *daGAL4>hsr*ω*RNAi*, which would have primarily affected the long nuclear transcripts containing the 280bp tandem repeats (Mallik and Lakhotia, 2009), showed milder defects in oogenesis than in *hsr*ω*^66^* ovaries, we selected *hsr*ω*RH* transcripts for ectopic expression. Remarkably, while expression of this transgene with either *ActGAL4* or *tjGAL4* driver in *hsr*ω*^66^* background resulted in partial improvement in oogenesis, its stronger and wider expression with a combination of *ActGAL4* and *tjGAL4* drivers in *hsr*ω*^66^* females substantially restored oogenesis, as revealed by the improved ovary morphology in *ActGAL4/tjGAL4>UAS hsr*ω*RH hsr*ω*^66^* flies and by the Caz and F-actin expression. Since the *hsr*ω*RH* transcript includes the region present in the *hsr*ω*RC* transcript and that encoding small Omega peptide (Sahu et al., 2021), the specific contributions of the two transcripts and Omega peptide in restoring oogenesis defects in *hsr*ω*^66^* ovaries remain unclear. This approach of utilizing the available GAL4-inducible transgenes for specific expression of specific transcripts of this gene (Sahu et al., 2021) provides directions for future studies to explore roles of each *hsr*ω transcript and the small Omega peptide in oogenesis in *Drosophila*.

The *hsr*ω lncRNA gene of *Drosophila melanogaster* has multi-faceted roles in normal development and in cell stress conditions (Lakhotia, 2011; Ray et al., 2019a; Ray et al., 2019b; Singh and Lakhotia, 2015). The present study extends this gene’s involvement in oogenesis at multiple stages via intra- as well as inter-organ regulatory events. As expected of a gene producing multiple transcripts which interact with a wide variety of regulatory and other molecules in cell, the present gene interaction studies did not reveal simple linear causal inter-relations with other potential regulators of oogenesis in *Drosophila.* These results nevertheless open novel and exciting possibilities for understanding roles of *hsr*ω and other lncRNAs in the complex process of oogenesis.

## Acknowledgements

We thank Pralay Majumder (Kolkata, India), Girish Ratnaparkhi (Pune, India) and the Bloomington Drosophila Stock Centre (USA) for providing fly stocks as listed in Materials and Methods.

## Materials and Methods

*Fly strains and genotypes*: All *Drosophila melanogaster* cultures were reared at 24^0^±1^0^C on standard food containing agar, maize powder, yeast, and sugar. Following genotypes were used: 1) *w^1118^* (as Control), 2) *w^1118^; hsr*ω*^66^* (Johnson et al., 2011), 3) *w^1118^; UAS-hsr*ω*-RNAi^3^*, targeting the 280pb tandem repeats in long *hsr*ω*RB hsr*ω*RG,* and *hsr*ω*RF* transcripts (Mallik and Lakhotia, 2009), and referred to here as *hsr*ω*RNAi*, 4) *w^1118^; UAS-hsr*ω*-exon2A-RNAi* targeting exon 2 in *hsr*ω lncRNAs (Sahu et al., 2021), and referred to here as *hsr*ω*exRNAi*, 5) *w^1118^; EP93D/TM6B* (Sengupta and Lakhotia, 2006) used for driving *EP93D* allele with GAL4 to over-express *hsr*ω transcripts, 6) *w*; daGAL4/TM6B* (BDSC no. 55851), a ubiquitous GAL4 driver (Chaturvedi et al., 2016), 7) *^w*^; tjGAL4/CyO*, a follicle cell specific driver (Weaver et al., 2020) received from Dr. Pralay Majumder (Presidency University, India), 8) *w*; Act5CGAL4/CyO-GFP* (BDSC no. 4414), a ubiquitous driver (Hongay and Orr Weaver, 2011) referred to here as *ActGAL4*, 9) *w^1118^; UAS-hsr*ω*-RH*, carrying the *UAS-hsr*ω *RH* transgene (Sahu et al., 2021) insertion on chromosome 3, referred to here as *hsr*ω*RH*, permitting targeted expression of the *hsr*ω-*RH* transcript, 10) *w*; UAS-cutRNAi* (BDSC no. 33967), referred to here as *cutRNAi*, 11) *w*; UAS-cut/CyO* (Yadav et al., 2022), used for over-expression of wild type Cut and referred to here as *cut*, 12) *UAS-Notch^act,^* for targeted expression of active Notch (Arya et al., 2015), referred to here as *N^act^*, 13) *y^1^ sc*^1^ v^1^ sev^21^; UAS-TBPH-RNAi/CyO; +/+* (BDSC no. 39014 (Lanson et al., 2011)), referred to here as *TBPHRNAi,* 14) *w^1118^; UAS-TBPHGFP* (Estes et al., 2013) expresses *TBPH* transcripts under UAS control, referred to here as TBPH. 15) *y^1^ sc*^1^ v^1^ sev^21^; +/+; UAS-cazRNAi* (dFus) (BDSC no. 34839), referred to here as *cazRNAi*, and 16) *w^1118^; +/+; UAS-FUS^WT^*, expresses wild type human *FUS*, a homolog of *Drosophila* Caz transcripts (Mallik et al., 2018) under UAS control (Lanson et al., 2011); referred to here as Fus,

Appropriate crosses were made to obtain progeny of desired genotypes.

### Adult Life-span assay

Progeny flies of desired genotypes were collected immediately after eclosion and grown in bottles/vials with food, with additional yeast for healthy growth. Three replicates of 20 females and males each of the desired genotypes were placed in food vials and transferred to fresh food vials at 3 days intervals till any *hsr*ω*^66^* female survived (maximum 33 days); following the transfer of flies to fresh vial, dead flies in the previous vial were examined for their sex and genotype.

### Fecundity assay

Flies of desired genotypes were collected soon after eclosion. Three replicates of 20 females and males each of the desired genotypes were placed in food vials as above and transferred into fresh food vials every day till day 14. Eggs laid in the previous vials during the one-day interval were counted to assay fecundity of females, which was expressed as eggs laid per female on a given day after eclosion.

### Measurement of 20-hydroxy-ecdysone (20-HE) levels in pupae and adults

30hr old *w^1118^* and *hsr*ω*^66^* female pupae and 4 days old adult females were used for measurement of 20-hydroxy ecdysone (20-HE) using the ecdysone immunoassay kit (Cat no. #A05120 Bertin Bioreagent, India) and following the manufacturer’s instructions. Three biological replicates were carried out, and a total of 150 female pupae and 114 adult females of each genotype were examined.

### 20-HE feeding

A stock solution (10mg/ml of ethanol) of 20-hydroxy-ecdysone (Cat no. H 5142 Sigma-Aldrich, India) was used to prepare ecdysone supplemented fly food with 20-HE at 0.05mM final concentration. The *w^1118^* and *hsr*ω*^66^* flies were collected immediately after eclosion and transferred to fresh regular food vials with additional yeast. On day 3 post eclosion, 8-9 females and 1 male of the given genotype (*w^1118^* or *hsr*ω*^66^*) were transferred to freshly prepared 20-HE supplemented food vials and allowed to feed for 115hr following which the 8 days old flies were anaesthetized and dissected to examine ovaries. Three biological replicates were carried out, and a total of 50 ovaries from 25 females of each genotype were examined.

### Immunostaining of ovaries

Ovaries were removed from 6 days old females of the desired genotypes, fixed, and processed for immunostaining as described earlier (Prasanth et al., 2000). The primary antibodies used were: 1) rat monoclonal anti-Vasa (1:10, DSHB), 2) mouse monoclonal anti-Alpha Spectrin (1:10, DSHB 3A9), 3) mouse monoclonal anti Fasciclin III (1:20, DSHB 7G10), 4) mouse monoclonal anti-Cut (1:10, DSHB 2B10), 5) mouse monoclonal Anti-Hnt (1:10, DSHB), and 6) mouse monoclonal Anti-Caz/dFus (1:10, 3F4) (Mallik et al., 2018), provided by Dr. Moushami Mallik (Germany)). Appropriate secondary antibodies conjugated with Alexa Flour 488 (1:200, Molecular Probes, USA) were used to detect the given primary antibody. Following the immunostaining, ovaries were counterstained with DAPI (4’,6-diamidino-2-phenylindole dihydrochloride, Sigma-Aldrich, India; 1µg/ml), washed thrice in PBST for 20 minutes each, mounted in DABCO (1,4-diazabicyclo[2.2.2]octane), and observed by fluorescence and/or confocal microscopy. At least 20 ovaries were immunostained and examined for each antibody and genotype.

### Phalloidin staining

Ovaries, fixed as above, were incubated in Phalloidin-TRITC (1:100, Sigma-Aldrich, India) for 12-16 hr at 4^0^C to detect filamentous actin (F-actin). Subsequent washing and counterstaining steps were as for immunostaining.

### Microscopy, quantification of signals and documentation

Confocal microscopy was carried out with Zeiss LSM 510 Meta laser scanning confocal microscope using Plan-Neofluor 5x (0.16 N.A.), Plan-Apo 10X (0.45 N.A.), Plan-Apo 20X (0.8 N.A.), or Plan-Apo 63X Oil (1.4 N.A.) with appropriate lasers and filters. Confocal images were analyzed with ImageJ software. For quantification of fluorescence signals in maximum intensity projection images, outlines of the germarium or other desired specific cells were marked with the “Roi manager” tool, and the signal intensity measured using the “Mean grey value” function in ImageJ. Width of germarium was measured using the “line tool” in ImageJ. All images were assembled with Adobe Photoshop 7.0. All confocal immunostaining images, unless otherwise stated, are presented as maximum intensity projection images while the DIC images for ovarian morphology (using Plan-Neofluor 5x objective) are presented as middle optical sections.

### Statistical analysis

Graph Pad Prism 8.4.2 was used for preparation of histograms and statistical analysis. For comparing the control and experimental sample values, Unpaired t test was applied. The significance of difference between the compared pairs of data is indicated in different histogram panels in the figures as: ns = not significant, *= p<0.05, **= p<0.01, ***= p<0.001 and ****= p<0.0001 above the corresponding bars.

