## Supplemental Material for "The multiple lncRNAs encoding *hsr*ω gene is essential for oogenesis in *Drosophila*"

**The *hsrω^66^* germaria show reduced Vasa and poorly developed fusomes**

The Terminal Filament and germ stem cell niche (Fig. 2A-B) and the total numbers of cap and terminal filament cells (Fig. 2E, N = 20 ovaries in each case) were comparable in *w^1118^* and *hsrω^66^* females, except the mid region of the germarium (region 2b) in *hsrω^66^* which appeared somewhat disorganized (Fig. 2D) and narrower (Fig. F) than in *w^1118^* (Fig. 2C).

Compared to *w^1118^* ovarioles (Fig. 2H), the germline marker RNA binding protein Vasa (Song et al., 2004; Yuan et al., 2012) was significantly weaker in 6 days old *hsrω^66^* germaria (Fig. 3H, I) although the mean numbers of the Vasa-positive GSCs (marked with ‘#’ in Fig. 2G, H) in *w^1118^* and *hsrω^66^* were comparable (Mean =2.06±0.01 and 2.02±0.01, respectively; N= 20 ovaries each).

The endoplasmic reticulum derived membranous fusomes grow along the cleavage furrows of cystoblasts with remnants of the mitotic spindle to connect and regulate the characteristic divisions of cystocytes during egg chamber differentiation (Greenbaum et al., 2011; Snapp et al., 2004). Immunostaining for α-Spectrin, one of the components of Fusomes, revealed (Fig. 2J-K) that unlike the branched and robust fusomes in *w^1118^* ovarioles (Fig. 2J), those in *hsrω^66^* were smaller, unbranched (Fig. 2K) and with significantly reduced α-Spectrin content (Fig. 2L). A detailed comparison of the distribution of α-Spectrin in later stage *w^1118^* and *hsrω^66^* could not be made because of their rarity in 6 days old *hsrω^66^* ovaries.

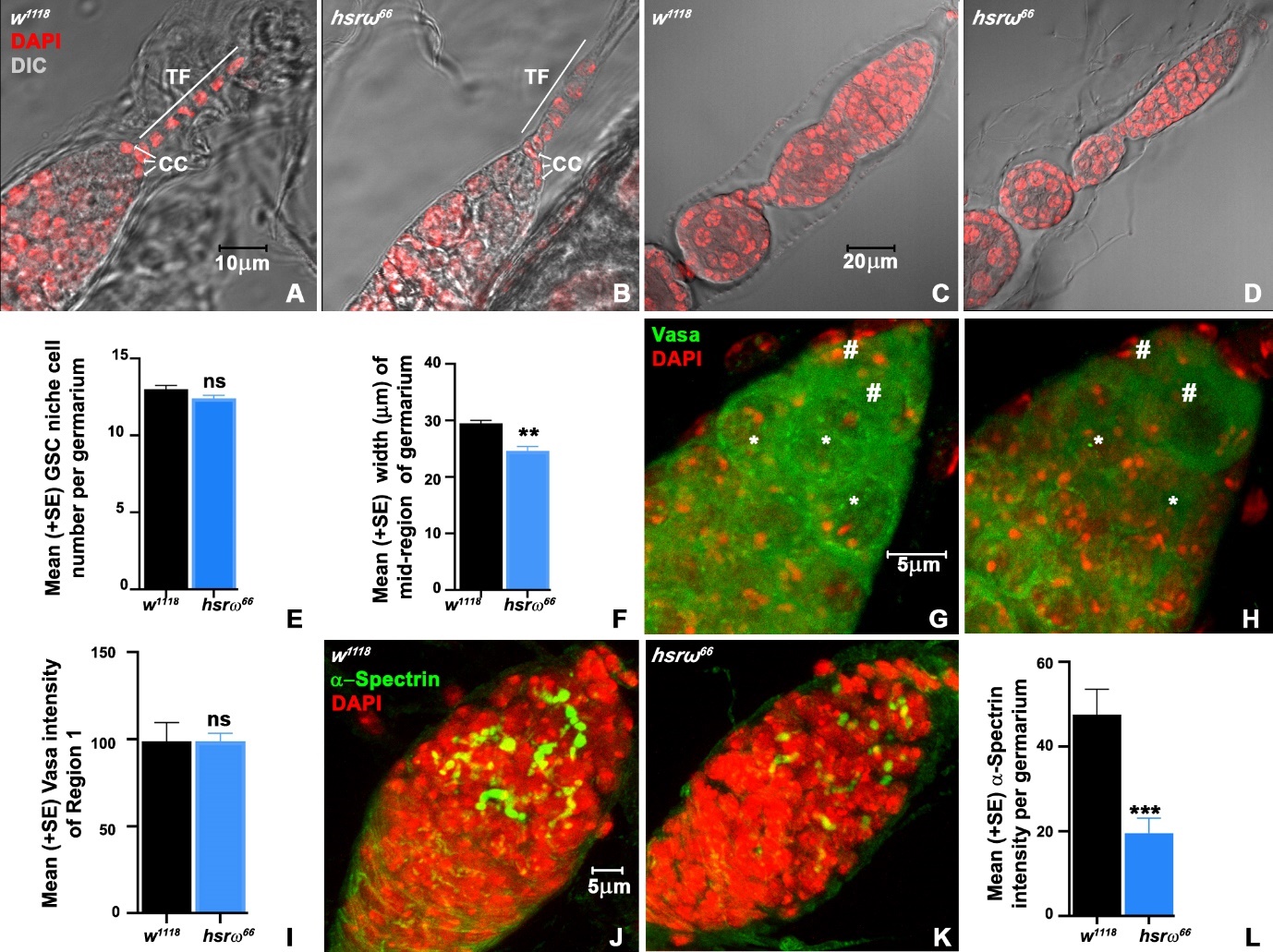

**Supplementary Fig. S1.** **Reduced Vasa and poorly formed fusomes in *hsrω^66^* ovaries.** **(A-D)** Germarium regions (DIC and DAPI stained, red) of 6 days old *w^1118^* (**A, C**) and *hsrω^66^* (**B, D**) females; white line and white arrows mark terminal filament cells (TF) and cap cells (CC), respectively. (**E-F**) **M**ean numbers of GSC (**E**, N= 10 for each genotype) cells and mean width of mid-region of germarium (**F**, N= 5 and 14 for *w^1118^* and *hsrω^66^*, respectively). (**G-H**) Confocal projection images showing Vasa (green) in germarium of 6 days old *w^1118^* (**G**) and *hsrω^66^* (**H**) females (DAPI, red); # and * mark GSCs and cystoblasts, respectively. (**I**) Vasa fluorescence intensities (in arbitrary units, Y-axis, N= 7 for each genotype) in region 1 of *w^1118^* and *hsrω^66^* ovarioles (X-axis). (**J, K**) α-Spectrin (green) distribution in anterior parts of *w^1118^* (**J**) and *hsrω^66^* (**K**) ovaries. (**L**) α-Spectrin fluorescence intensities (in arbitrary units, Y-axis, N= 8 and 18 for *w^1118^* and *hsrω^66^*, respectively) per germarium (genotypes on X-axis) Scale bar in **A** (10µm) applies to **A-B,** in **C** (20µm) to **C-D**, in **J** (5µm) to **J-K**.

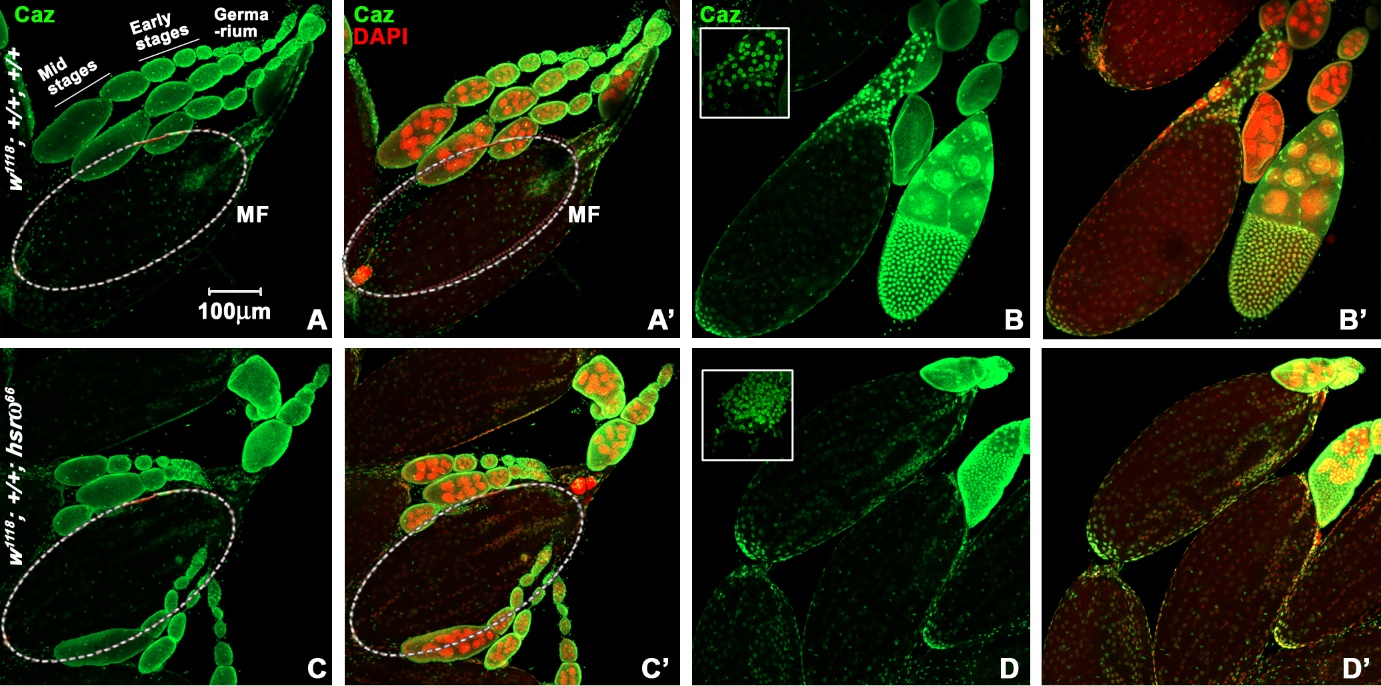

**Supplementary** **Fig. S2. Caz/dFus hnRNP expression in *w^1118^* and *hsrω^66^* ovarioles generally similar but levels reduced specially in mature *hsrω^66^* follicles. (A-D**) of Caz (green) and DAPI (red, **A’**-**D’**) stained ovarioles from 6 days old *w^1118^* (**A,** **B**) and *hsrω^66^* (**C,** **D**) females; insets in **B** and **D** show Caz (green) in anterior follicle cells at early stage 14 chambers. Germarium, early- and mid-stage follicles are marked in **A;** mature follicles are outlined in **A**, **A’**, **C**, and **C’**. Scale bar in **A** (100µm) applies to **A**-**D’**.
